# Comparison of the *in vitro*-efficacy of different mouthwash solutions targeting SARS-CoV-2 based on the European Standard EN 14476

**DOI:** 10.1101/2020.10.25.354571

**Authors:** Katrin Steinhauer, Toni Luise Meister, Daniel Todt, Adalbert Krawczyk, Lars Paßvogel, Britta Becker, Dajana Paulmann, Birte Bischoff, Stephanie Pfaender, Florian H H Brill, Eike Steinmann

## Abstract

The SARS-Cov-2 pandemic is triggering a global health emergency alert, and recent research is indicating the relevance of aerosols in the spread of SARS-CoV-2. Thus, in this study antiseptic mouthwashes based on the actives chlorhexidine (CHX) and octenidine (OCT) were investigated regarding their efficacy against SARS-CoV-2 using EN 14476. Based on the requirement of EN 14476 (i.e. reduction of viral titer by ≥ 4 log _10_), the OCT-based formulation was effective within only 15 sec against SARS-CoV-2, and thus constitutes an interesting candidate for future clinical studies to prove its effectiveness in a potential prevention of SARS-CoV-2 transmission by aerosols.

## Introduction

Coronaviruses are enveloped single-stranded RNA viruses and are characterized by club shaped spikes on the surface of the virion, prompting the name coronavirus due to the similarity in appearance to a solar corona [1]. Until the SARS-CoV outbreak in 2002, coronaviruses were thought to only cause mild self-limiting infections in humans but were known to cause a wide variety of infections in animals [1]. 17 years later, in December 2019, a novel coronavirus was identified as the causative agent of severe pneumonia in a cluster of patients [2], designated as SARS-CoV-2 due to its relatedness to severe acute respiratory syndrome coronavirus (SARS-CoV) [3]. Since then SARS-CoV-2 spread around the world thereby triggering a global health emergency alert. Thus, until vaccination becomes available a bundle of effective preventive measures is desperately needed.

In this context, recent publications suggest the use of antimicrobial mouthwashes as a preventive measure. This is based on the efficacy of antimicrobial mouthwashes to reduce the number of microorganisms in the oral cavity prompting a reduction of microorganisms in aerosols [4]. This is particularly interesting, as recent research indicates the relevance of aerosols also in the spread of SARS CoV-2 [5].

Thus, in their review summarizing data for mouthwashes with chlorhexidine gluconate (CHX), cetylpyridinium chloride (CPC), povidone-iodine (PVP-I), and hydrogen peroxide (H_2_O_2_) Vergara-Beunaventura and Castro-Ruiz indicate an essential role of antiseptic mouthwashes to reduce SARS-CoV-2 viral load in dental practice and undermine that research on this topic is urgently needed to verify the potential of antiseptic mouth rinses as a further preventive measure [6]. The aim of our study was therefore, to directly compare commercially available antiseptic mouthwash formulations based on the common antiseptic actives chlorhexidine (CHX) and octenidine dihydrochloride (OCT) regarding their efficacy against the pandemic coronavirus SARS-CoV-2. *In vitro*-experiments were carried out in accordance to the well-established European Standard EN 14476 [7] determined for evaluating the virucidal efficacy of chemical disinfectants and antiseptics, in which reduction of at least four decimal logarithms (log_10_) of viral titer is requested to state efficacy.

## Material and Methods

### Quantitative Suspension tests according to EN 14476

Quantitative suspensions tests were carried out as described in EN 14476 [7]. Briefly, efficacy against SARS CoV-2 [8] was studied using commercially available mouthwashes. A commercially available ready-to-use formulation designated formulation A (100 g contains: 0.1 g Chlorhexidine bis-(D-gluconate); GlaxoSmithKline Consumer Health GmbH & Co. KG, Germany) was used as one test formulation. In addition, a commercially available ready-to-use formulation designated formulation B (100 g contains: 0.2 g Chlorhexidine bis-(D-gluconate); GlaxoSmithKline Consumer Health GmbH & Co. KG, Germany) was used. Formulation C used in this study was also a ready-to-use preparation (trade name: octenisept, (drug authorisation number: 32834.00.00); 100 g contains: 0.1 g octenidine dihydrochloride, 2 g phenoxyethanol). Concentrations and contact times used throughout this study are indicated. Experiments were carried out under conditions of low organic soiling (0.3 g/L bovine serum albumin (BSA); “clean conditions”) as defined in EN 14476 [7].

Data presented are based on at least two independent experiments. Validation controls as defined in EN 14476 [7] were found to be effective in all experiments indicating validity of presented data.

## Results and Discussion

Comparison of different commercially available mouthwashes based on the well-established antiseptically effective actives chlorhexidinedigluconate (CHX) and octenidinedihydrochloride (OCT) was conducted considering the European Standard EN 14476 [7]. The assays were carried out using an isolated SARS-CoV-2 outbreak strain [8] in the presence of low organic soiling (i.e. 0.3 g/L bovine serum albumin) as requested by EN 14476 [7] to also consider potential quenching of the actives by protein load as has been described before [9].

Data is presented in Figure 1. Figure 1 A shows SARS-CoV-2 reduction obtained for products A, B and C using end point titration. In these experiments the two formulations based on chlorhexidinedigluconate (formulations A and B) were found to have only limited efficacy against SARS-CoV-2. Thus, at a concentration of 80% (v/v) formulation A containing 0.1 % chlorhexidinedigluconate reduced the virus titer even at a prolonged contact time of 10 min by less than 1 log_10_. Formulation B containing 0.2 % chlorhexidinedigluconate reduced SARS-CoV-2 within a contact time of 1 min as well as at a prolonged contact time of 5 min when tested at 80% (v/v) concentration also by less than 1 log_10_. As for formulations A and B reduction of the virus titer was found not to be impacted by cytotoxic effects of the formulations, indicated by the lower limit of quantification (LLOQ), no additional large-volume plating (LVP) experiments were conducted. This is well in line with data from screening experiments in our lab, where virus reduction titers were found not to be elevated due to less toxicity when using both formulations at a concentration of only 20% (v/v) (data not shown).

**Figure 1:**
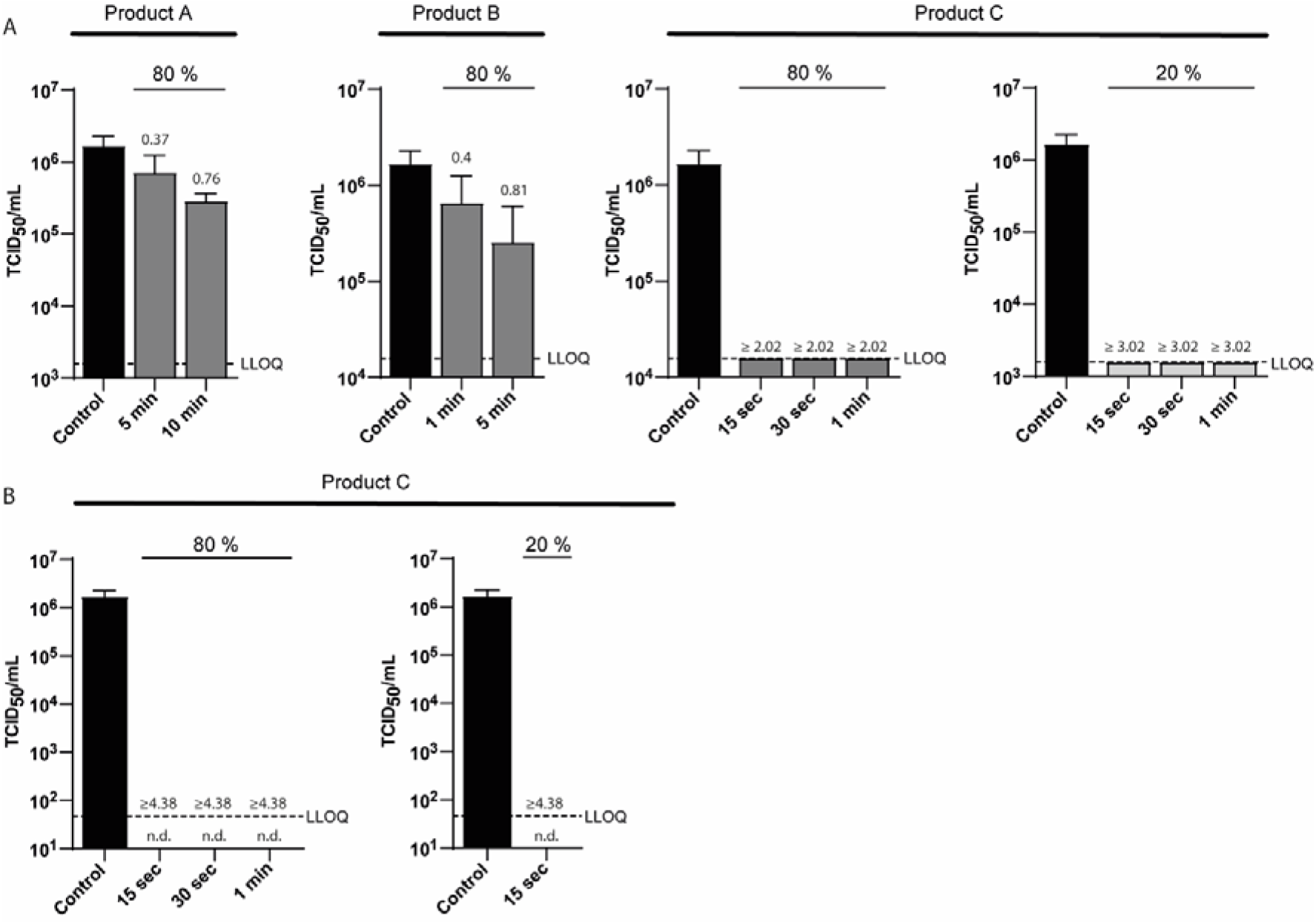
Virucidal activity of oral rinses against SARS-CoV-2. SARS-CoV-2 was incubated with medium (control, black bar) or various oral rinses (Product A-C) for indicated concentrations (80 % and/or 20 %) and time periods (15 sec to 10 min). The cytotoxic effect was monitored using non-infected cells incubated with the different products, defined as lower limit of quantification (LLOQ). Log-reduction factors are indicated above the bars. In panal A viral titers were determined upon limited endpoint titration on Vero E6 cells. Tissue culture infectious dose 50 (TCID50/mL) was calculated according to Spearman-Kärber. Due to high cytotoxic effects diminishing the measuring window for product C large volume plating was performed to reduce cytotoxicity and evaluate the remaining titers below 10^4 (panel B). No remaining cytopathic effects were observed (n.d.).

In contrast, when looking at the data for formulation C logarithmic reduction factors log_10_ were found to be 1 log_10_ higher (i.e. ≥ 3.02 log_10_) for the 20% (v/v) concentration of product C when compared to the 80% (v/v) test concentration (i.e. ≥ 2.02 log_10_). This indicates, that the measuring window for product was diminished by cytotoxicity. Therefore, additional large volume plating (LVP) experiments to obtain a wider measuring window were conducted with formulation C. Data obtained using LVP are presented in figure 1 B, and indicate a reduction of SARS-CoV-2 titers by ≥ 4.38 log_10_ already within the shortest contact time of 15 sec for the octenidine dihydrochloride (OCT)-based mouthwash (formulation C). This was found for both test concentrations tested (80% (v/v) and 20% (v/v)).

Data presented in this study for the two CHX-based mouthwashes (formulations A and B) are well in line with data published by Meister et al. [10]. In their investigation of different mouthwashes targeting SARS-CoV-2 also only a limited efficacy (i.e. < 1 log_10_) of the two tested commercially available mouthwashes based on chlorhexidinedigluconate was found. However, looking at the data for the octenidinedihydrochloride-based mouthwashes, in the earlier study by Meister et. al. [10] only limited virucidal activity of the formulation tested (i.e. < 1 log_10_) was found, whereas in this study the tested octenidinedihydrochloride-based formulation (C) was found effective against SARS-CoV-2 within 15 sec (i.e. ≥ 4 log_10_). This differing data can be explained by the use of two different octenidine-based formulations in the two studies. In the earlier study [10] a formulation containing octenidinedihydrochloride as the only active was used as compared to the OCT-based formulation (formulation C) used in this study which contained octenidinedihydrochloride in combination with phenoxyethanol. Furthermore this discrepancy indicates the value of pre-evaluating each individual formulation on the basis of EN 14476 when assessing the virucidal potential against SARS CoV-2. For this pre-evaluation the standard test surrogate virus modified vaccinia virus strain Ankara (MVA) to assess “virucidal activity against enveloped viruses” as defined in EN 14476 [7] has been found to be of value, as with this approach a non-pathogenic virus can be used in the lab to obtain reliable data regarding virucidal activity against enveloped viruses in general including SARS CoV-2.

In conclusion, in this study virucidal efficacy against SARS-CoV-2 could be demonstrated for formulation C meeting the > 4 log_10_ requirement of EN 14476 [7] within a contact time of only 15 sec, making this formulation suitable to be used as a mouthwash.

Thus, based on this *in vitro*-data the OCT-based formulation used in this study constitutes an interesting candidate for future clinical studies to prove its effectiveness in a potential prevention of SARS-CoV-2 spread by aerosols.

